# Detection of SARS-CoV-2 RNA by multiplex RT-qPCR

**DOI:** 10.1101/2020.06.16.155887

**Authors:** Eriko Kudo, Benjamin Israelow, Chantal B.F. Vogels, Peiwen Lu, Anne L. Wyllie, Maria Tokuyama, Arvind Venkataraman, Doug E Brackney, Isabel M. Ott, Mary E. Petrone, Rebecca Earnest, Sarah Lapidus, M. Catherine Muenker, Adam J. Moore, Arnau Casanovas-Massana, Yale IMPACT Research Team, Saad B. Omer, Charles S. Dela Cruz, Shelli F. Farhadian, Albert I. Ko, Nathan D. Grubaugh, Akiko Iwasaki

**Affiliations:** Department of Immunobiology, Yale University School of Medicine, New Haven, CT, 06520, USA; Department of Internal Medicine, Section of Infectious Diseases, Yale School of Medicine, New Haven, CT, 06510, USA; Department of Epidemiology of Microbial Diseases, Yale School of Public Health, New Haven, CT 06510, USA; Yale Institute of Global Health, New Haven, CT 06510, USA; Yale School of Nursing, New Haven, CT 06510, USA; Department of Internal Medicine, Section of Pulmonary, Critical Care, and Sleep Medicine, Yale School of Medicine, New Haven, CT, 06510, USA; Howard Hughes Medical Institute, Chevy Chase, MD, 20815, USA

**Keywords:** SARS-CoV-2, COVID-19, multiplex RT-qPCR

## Abstract

The current RT-qPCR assay recommended for SARS-CoV-2 testing in the United
States requires analysis of three genomic targets per sample: two viral and one host. To simplify testing and reduce the volume of required reagents, we developed a multiplex RT-qPCR assay to detect SARS-CoV-2 in a single reaction. We used existing N1, N2, and RP primer and probe sets by the CDC, but substituted fluorophores to allow multiplexing of the assay. The cycle threshold (Ct) values of our multiplex RT-qPCR were comparable to those obtained by the singleplex assay adapted for research purposes. Low copies (>500 copies / reaction) of SARS-CoV-2 RNA were consistently detected by the multiplex RT-qPCR. Our novel multiplex RT-qPCR improves upon current singleplex diagnostics by saving reagents, costs, time and labor.

## Introduction

The ongoing global pandemic caused by severe acute respiratory syndrome coronavirus 2 (SARS-CoV-2) and associated coronavirus disease 2019 (COVID-19) has caused more than 7.5 million infections and killed more than 423,000 people as of June 14, 2020, and the virus continues to spread throughout the globe [1]. In the absence of a specific vaccine or effective therapy for the treatment of COVID-19, public health infection prevention measures, including contact tracing and isolation measures, are currently our only tool to stem transmission. However, testing, contact tracing and isolation measures require rapid and widespread testing. Here, we developed a quantitative reverse transcription PCR (RT-qPCR) assay for the detection of SARS-CoV-2 to allow for more rapid and widespread testing.

While a number of primer and probe sets for the detection of SARS-CoV-2 RNA by RT-qPCR have become available since the identification of this novel virus, its broad deployment has been hampered partially by the availability of testing reagents. The current RT-qPCR assay developed by the CDC targets two different conserved segments of the viral nucleocapsid gene (N1 and N2) as well as the human RNase P gene as a sampling control [2]. This protocol therefore requires 3 reactions to be performed per patient sample, which, in addition to requiring a large amount of resources, also increases the chance for error. In an effort to reduce reagents, time, potential error and labor per sample, we developed a multiplex RT-qPCR for the detection of SARS-CoV-2. To do this, we utilized the existing N1 and N2 primer and probe sets published by the CDC; however, we substituted different fluorophores to enable multiplexing. We found the accuracy and specificity of this method to be similar to singleplex RT-qPCR. While there are commercially available tests that employ multiplex PCR, their methods remain proprietary to the companies and are not published. Important consideration in this regard is the such tests are cost prohibitive in low and middle income countries in which COVID-19 pandemic is spreading. Therefore, this novel multiplex RT-qPCR assay provides the first publicly available multiplex PCR protocol, which provides equivalent diagnostic accuracy to current singleplex methods in fewer reactions and utilizes less reagents and time.

## Materials and Methods

### Clinical samples

Clinical samples from Yale-New Haven Hospital COVID-19 diagnosed inpatients and health care workers were collected as part of Yale’s project IMPACT biorepository. RNA was extracted from nasopharyngeal and saliva samples using the MagMax Viral/Pathogen Nucleic Acid Isolation kit (ThermoFisher Scientific, Waltham, MA, USA), according to a modified protocol [3].

### Control samples

Full-length SARS-CoV-2 RNA (WA1_USA strain from University of Texas Medical Branch (UTMB); GenBank: MN985325) [4] was used as positive control for validation. Total RNA extracted from human embryonic kidney cell line 293T was used for detection of internal host gene control.

### Singleplex and multiplex RT-qPCR

All reactions were performed on a CFX96 Touch instrument (Bio-Rad, Hercules, CA, USA) using Luna Universal Probe One-Step RT-qPCR Kit (New England BioLabs, Ipswich, MA, USA) according to the manufacturer’s protocol. A final reaction volume of 20 μl containing 5 μl template was used. The following cycling conditions were applied; a cDNA synthesis 10 min/55°C, a hold step 1 min/95°C, and subsequently 45 cycles of denaturation 10 s/95°C and annealing/elongation 30 s/55°C. Nuclease-free water was used as the non-template control. The primer pairs and probes for single- and multiplex RT-qPCR are shown in **Table 1**. We calculated analytic efficiency of RT-qPCR assays tested with full-length SARS-CoV-2 RNA using the following formula.

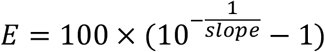

**Table 1.**
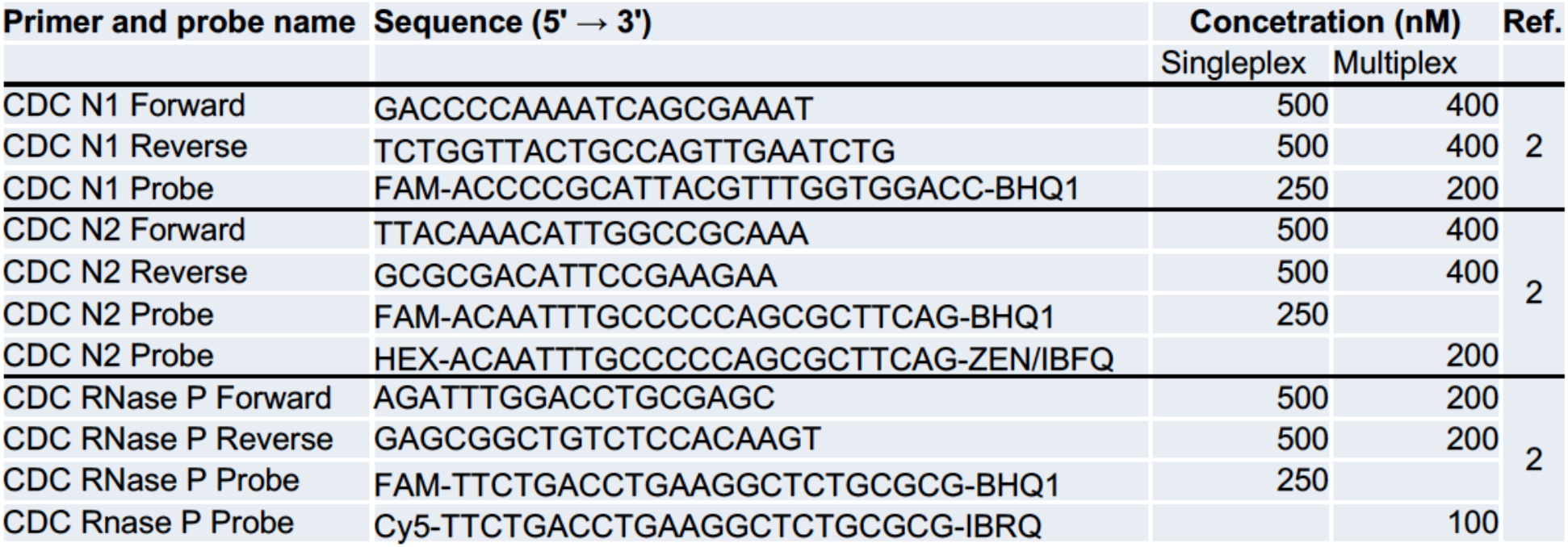
Primer and probe for single- and multiplex RT-qPCR

## Results

### Determination of lower limit of virus concentration detected by multiplex RT-qPCR

The limit of detection (LOD) was analyzed using 10-fold serially diluted full-length SARS-CoV-2 RNA into RNA extracted from pooled nasopharyngeal swabs from COVID-uninfected research participants. The Ct values and detection rates are shown in **Table 2**. The slope of the standard curves for N1 and N2 were −3.36 and −3.52, respectively. The amplification efficiency was above 90% for both primer probe sets (**Figure 1**). All primer-probe sets and conditions were able to detect SARS-CoV-2 at 500 virus copies per reaction **(Table 2)**.

**Table 2.** Lower limit of detection of SARS-CoV-2 in multiplex RT-qPCR

| Copies/reaction | Ct: average (SD) |  | Detection/tested (%) |  |
| --- | --- | --- | --- | --- |
|  | N1 | N2 | N1 | N2 |
| 5 | 39.24 (0.38) | 39.63 | 3/20 (15) | 1/20 (5) |
| 50 | 38.59 (0.64) | 38.51 (0.84) | 12/20 (60) | 5/20 (25) |
| 500 | 36.50 (0.63) | 36.67 (0.87) | 20/20 (100) | 20/20 (100) |
| 5000 | 32.56 (0.30) | 32.20 (0.27) | 20/20 (100) | 20/20 (100) |
| 50000 | 28.70 (0.22) | 28.23 (0.17) | 20/20 (100) | 20/20 (100) |

**Fig. 1.**
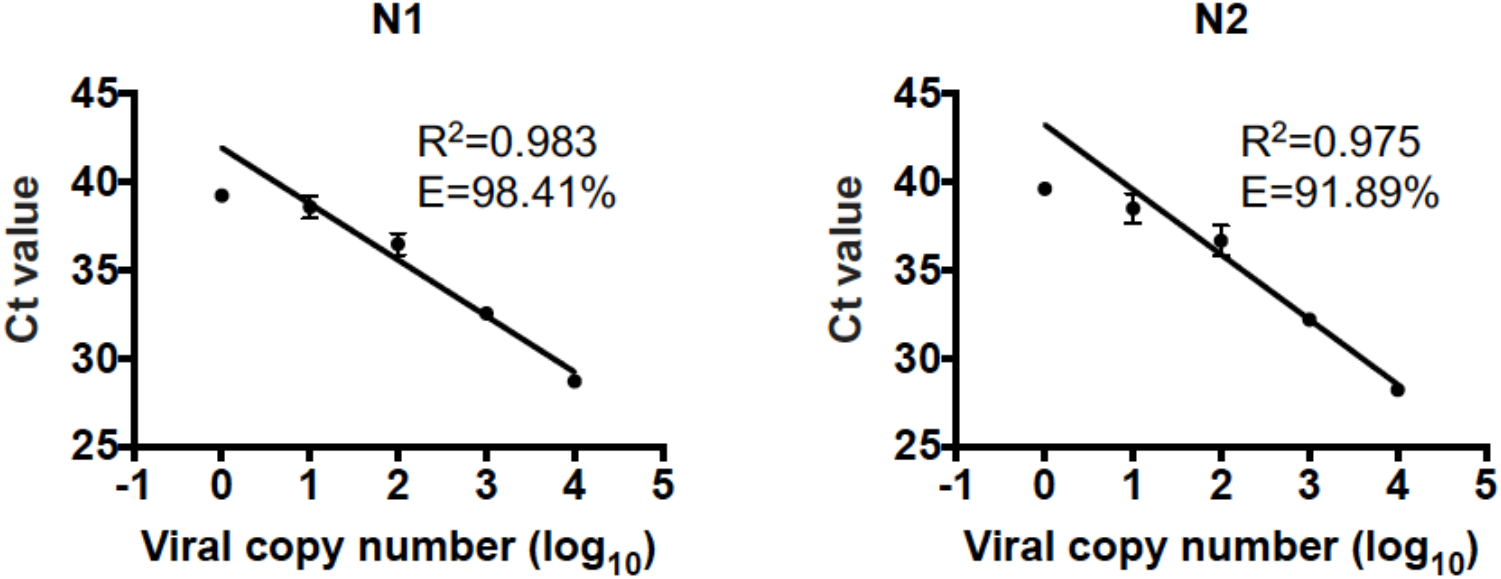
Standard curve generated for N1 and N2 of SARS-CoV-2. Multiplex RT-qPCR detection of SARS-CoV-2 N1 and N2 gene were validated using ten-fold dilutions of viral RNA into pooled negative nasopharyngeal swabs samples measure sensitivity and efficiency for twenty replicates. Data are mean ± SD. R^2^, regression coefficient value; E, amplification efficiency.

### Comparison of performance of multiplex and singleplex RT-qPCR

To confirm the specificity of the primer-probe sets (FAM, HEX, and Cy5 fluorophores) either tested as single or multiplex reactions, as well as in comparison to the original singleplex assay (FAM), we used nasopharyngeal swab and saliva samples from patients to detect SARS-CoV-2 RNA. The Ct values generated by the multiplex RT-qPCR were similar with FAM only or multi-color of singleplex RT-qPCR **(Figure 2 and Table 3)**. These data indicated that our RT-qPCR with multicolor fluorophores under single- and multiplex conditions has similar performance for the detection of SARS-CoV-2 RNA as the currently utilized singleplex RT-qPCR.

**Figure 2.**
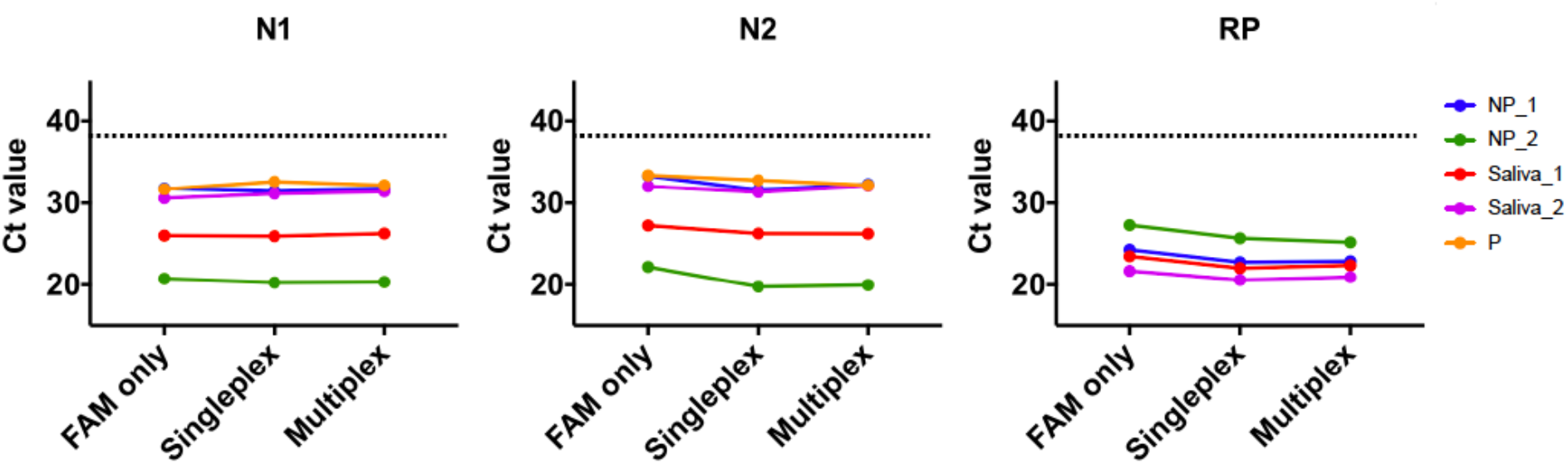
Results of Ct value in singleplex and multiplex RT-qPCR. Four independent COVID-19 inpatients’ nasopharyngeal (n=2) or saliva (n=2) samples, one negative control and one positive control (P) (10^3^ virus copy per μl) were compared to the Ct value between single- or multi-color singleplex RT-qPCR and multiplex RT-qPCR. Dash line indicates the cut-off value of 38 Ct. P, positive control. Negative control was undetectable. Individual values are indicated in Table 3.

**Table 3.** Comparison of Ct value between singleplex and multiplex RT-qPCR

| Sample | Singleplex (only FAM) |  |  | Singleplex |  |  | Multiplex |  |  |
| --- | --- | --- | --- | --- | --- | --- | --- | --- | --- |
|  | N1 | N2 | RP | N1 | N2 | RP | N1 | N2 | RP |
| NP_1 | 31.78 | 33.31 | 24.28 | 31.44 | 31.58 | 22.75 | 31.73 | 32.30 | 22.89 |
| NP_2 | 20.73 | 22.19 | 27.29 | 20.25 | 19.75 | 25.69 | 20.34 | 19.96 | 25.20 |
| Saliva_1 | 26.01 | 27.24 | 23.48 | 25.93 | 26.27 | 22.03 | 26.26 | 26.23 | 22.39 |
| Saliva_2 | 30.58 | 32.04 | 21.69 | 31.15 | 31.33 | 20.59 | 31.41 | 32.08 | 20.94 |
| P | 31.68 | 33.4 | ND | 32.60 | 32.80 | ND | 32.17 | 32.17 | ND |
| N | ND | ND | ND | ND | ND | ND | ND | ND | ND |

N, negative control; P, Positive control (SARS-CoV-2 RNA with a concentration of $10^3$ per $\mu\text{l}$ ); ND, not detected.

### Comparison of single- and multiplex assay sensitivity with clinical samples

To evaluate the accuracy of our RT-qPCR multiplex assay, we tested RNA extracted from nasopharyngeal swabs and saliva samples obtained from COVID-19-positive hospitalized patients and COVID-19-uninfected health care workers. Total of 42 samples included 34 COVID-19-positive inpatients and 8 uninfected health care workers. The results of our multiplex RT-qPCR were 100% sensitive as compared with singleplex RT-qPCR **(Figure 3 and Table 4)**. These data show that our multiplex RT-qPCR method could provide an alternative to the detection of SARS-CoV-2 by currently published singleplex methods.

**Figure 3.**
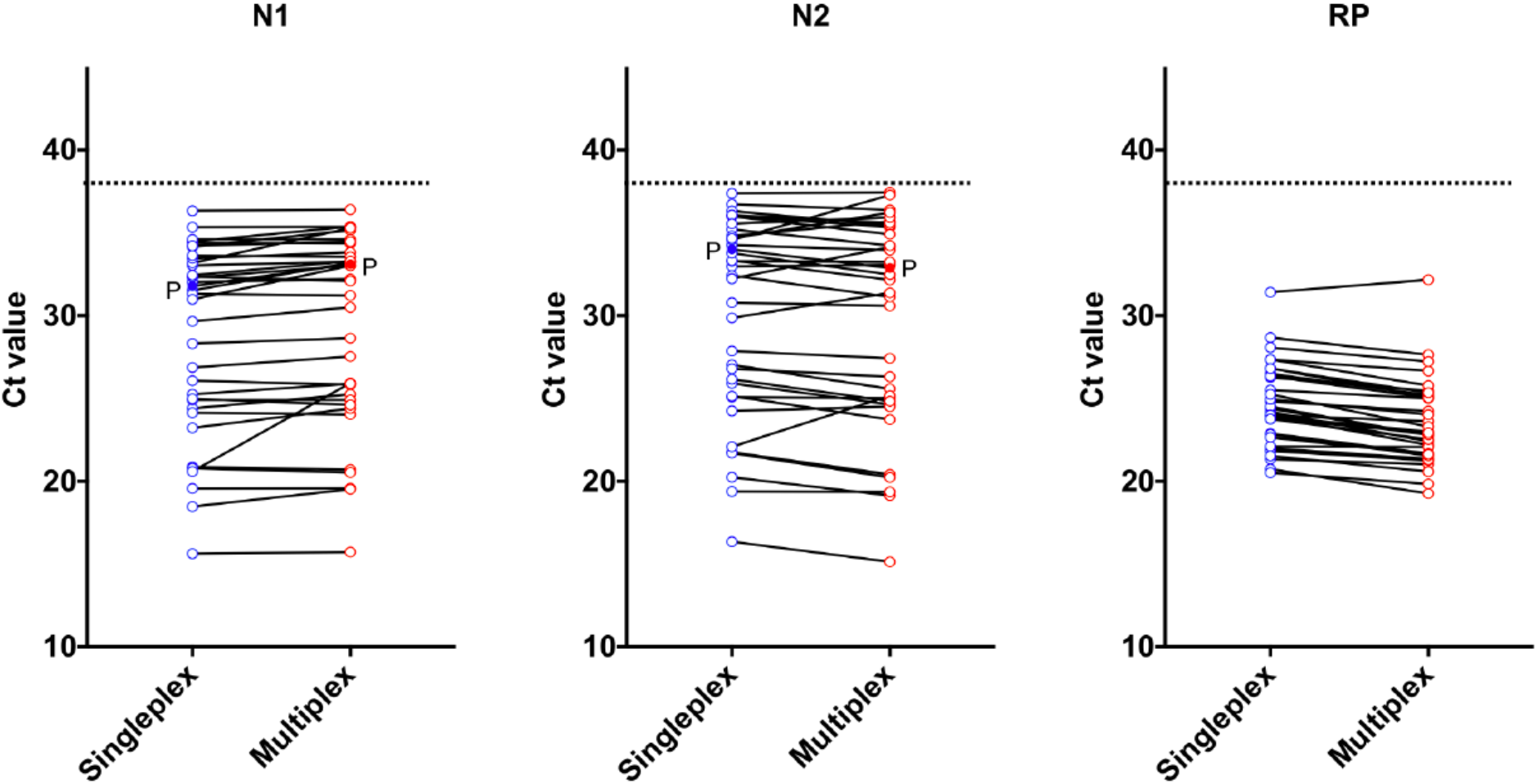
Comparison of single- and multiplex assay performance with clinical samples. Forty two RNA templates from nasopharyngeal swabs and saliva samples obtained from COVID-19 inpatients or health care worker and positive control (P) (10^3^ virus copy per μl) were performed singleplex and multiplex RT-qPCR. Dash line indicates the cut-off value of 38 Ct. Individual values are indicated in Table 4.

**Table 4.** The Ct value and result from multiplex assay in clinical samples

| Sample | Sample type | Singleplex |  |  |  | Multiplex |  |  |  |
| --- | --- | --- | --- | --- | --- | --- | --- | --- | --- |
|  |  | N1 | N2 | RP | Result | N1 | N2 | RP | Result |
| 1 | NP | 32.29 | 32.96 | 23.99 | Positive | 33.01 | 33.06 | 23.62 | Positive |
| 2 | NP | 29.65 | 30.78 | 25.51 | Positive | 30.50 | 30.58 | 25.00 | Positive |
| 3 | NP | 31.52 | 33.29 | 31.41 | Positive | 33.20 | 33.24 | 32.16 | Positive |
| 4 | NP | 24.4 | 25.07 | 24.82 | Positive | 25.24 | 25.02 | 24.23 | Positive |
| 5 | NP | 19.57 | 20.24 | 23.94 | Positive | 19.58 | 19.13 | 22.20 | Positive |
| 6 | NP | 35.33 | 36.72 | 23.93 | Positive | 35.36 | 36.37 | 22.98 | Positive |
| 7 | NP | 34.46 | 36.03 | 26.27 | Positive | 35.37 | 35.34 | 25.46 | Positive |
| 8 | NP | 24.13 | 25.13 | 26.49 | Positive | 24.03 | 23.73 | 25.20 | Positive |
| 9 | NP | 24.9 | 25.91 | 27.34 | Positive | 24.90 | 24.63 | 25.78 | Positive |
| 10 | NP | 31.31 | 32.41 | 28.67 | Positive | 31.21 | 31.12 | 27.64 | Positive |
| 11 | NP | 26.07 | 27.02 | 26.82 | Positive | 25.83 | 25.58 | 25.09 | Positive |
| 12 | NP | 34.15 | 36.31 | 27.34 | Positive | 35.15 | 35.45 | 26.65 | Positive |
| 13 | NP | 36.32 | 37.37 | 26.35 | Positive | 36.40 | 37.44 | 25.03 | Positive |
| 14 | NP | 32.03 | 33.32 | 26.8 | Positive | 32.20 | 32.15 | 25.35 | Positive |
| 15 | Saliva | 34.39 | 35.99 | 24.38 | Positive | 34.35 | 34.91 | 22.80 | Positive |
| 16 | Saliva | 32.34 | 33.77 | 24.19 | Positive | 32.09 | 32.47 | 22.41 | Positive |
| 17 | Saliva | 30.98 | 32.23 | 21.33 | Positive | 33.02 | 34.14 | 21.02 | Positive |
| 18 | Saliva | 15.63 | 16.36 | 22 | Positive | 15.73 | 15.13 | 21.36 | Positive |
| 19 | Saliva | 18.46 | 19.38 | 21.87 | Positive | 19.51 | 19.36 | 21.25 | Positive |
| 20 | Saliva | 26.86 | 27.86 | 21.82 | Positive | 27.53 | 27.42 | 21.29 | Positive |
| 21 | Saliva | 23.22 | 24.25 | 22.11 | Positive | 24.40 | 24.50 | 22.07 | Positive |
| 22 | Saliva | 20.86 | 21.75 | 24.5 | Positive | 20.70 | 20.41 | 22.77 | Positive |
| 23 | Saliva | 33.03 | 34.25 | 24.95 | Positive | 33.25 | 33.95 | 24.02 | Positive |
| 24 | Saliva | 20.77 | 21.7 | 25.25 | Positive | 20.53 | 20.23 | 23.29 | Positive |
| 25 | Saliva | 33.17 | 34.56 | 21.53 | Positive | 35.29 | 37.28 | 20.58 | Positive |
| 26 | Saliva | 33.46 | 35.2 | 28.07 | Positive | 33.82 | 34.23 | 27.22 | Positive |
| 27 | Saliva | 33.63 | 34.81 | 22.87 | Positive | 33.54 | 35.68 | 21.68 | Positive |
| 28 | Saliva | 20.59 | 22.09 | 22.83 | Positive | 25.92 | 25.16 | 21.54 | Positive |
| 29 | Saliva | 25.24 | 26.79 | 20.72 | Positive | 25.89 | 26.31 | 19.26 | Positive |
| 30 | Saliva | 34.6 | 36.07 | 23.75 | Positive | 34.59 | 35.60 | 22.53 | Positive |
| 31 | Saliva | 24.97 | 26.16 | 24 | Positive | 24.60 | 24.83 | 22.81 | Positive |
| 32 | Saliva | 34.21 | 35.55 | 23.77 | Positive | 34.46 | 35.93 | 22.92 | Positive |
| 33 | Saliva | 28.31 | 29.86 | 22.65 | Positive | 28.63 | 31.37 | 21.63 | Positive |
| 34 | Saliva | 32.43 | 34.66 | 20.51 | Positive | 33.29 | 36.23 | 19.84 | Positive |
| 35 | NP | ND | ND | 29.6 | Not detected | ND | ND | 28.78 | Not detected |
| 36 | NP | ND | ND | 27.03 | Not detected | ND | ND | 26.09 | Not detected |
| 37 | NP | ND | ND | 29.86 | Not detected | ND | ND | 29.24 | Not detected |
| 38 | NP | ND | ND | 29.34 | Not detected | ND | ND | 28.50 | Not detected |
| 39 | Saliva | ND | ND | 24.56 | Not detected | ND | ND | 23.15 | Not detected |
| 40 | Saliva | ND | ND | 25.35 | Not detected | ND | ND | 23.96 | Not detected |
| 41 | Saliva | ND | ND | 24.9 | Not detected | ND | ND | 23.63 | Not detected |
| 42 | Saliva | ND | ND | 26.38 | Not detected | ND | ND | 25.12 | Not detected |
| P |  | 31.78 | 34.00 | ND |  | 33.10 | 32.86 | ND |  |

RP, human RNase P; P, Positive control (SARS-CoV-2 RNA with a concentration of $10^3$ viral copies per $\mu\text{l}$ ); ND, not detected; NP, nasopharyngeal swab. (1,742)

## Discussion

We developed a multiplex RT-qPCR for molecular diagnostic testing of SARS-CoV-2 by improving on an existing research singleplex RT-qPCR method using the CDC primer-probe sets. This multiplex RT-qPCR approach simultaneously detected the CDC-recommended two gene segments of the SARS-CoV-2 (N1 and N2) and the internal control human RNase P gene in a single reaction for research purposes. This method performed as well as the singleplex RT-qPCR with clinical samples and was very specific for detecting all target genes. Generally, an important consideration for this multiplex RT-qPCR approach is that cycling conditions may vary depending on qPCR machines, sample type and target gene. We therefore recommend that when implementing new assays, primer and probe concentrations should be optimized to individual lab conditions.

The US CDC primer and probe sets for COVID-19 testing are recommended for clinical testing in the US [2]. We reported sensitivity of US CDC primer and probe sets compared with others; China CDC [5], Charité Institute of Virology, Universitätsmedizin Berlin [6] and Hong Kong University [7]. In singleplex RT-qPCR, CDC N2 primer set has a lower detection capability than CDC N1 primers [8]. Our multiplex RT-qPCR assay also showed that N1 and N2 primer-probe set were 60 % and 25 % detection in 50 copies per reaction, respectively (**Table 2**).

The SARS-CoV-2 pandemic has already claimed the lives of over 400,000 people, and halted the global economy and changed our daily lives worldwide. A rapid and accurate diagnosis that is not cost prohibitive to test for infected individuals is urgently needed. Our multiplex RT-qPCR protocol described in this study provides rapid and highly sensitive detection of SARS-CoV-2 RNA for research purposes. In the future, FDA approval of such multiplex PCR technique for clinical testing could provide a cost effective solution to mass testing.

## Yale IMPACT Research Team Authors

(Listed in alphabetical order) Kelly Anastasio, Michael H. Askenase, Maria Batsu, Santos Bermejo, Sean Bickerton, Kristina Brower, Melissa Campbell, Yiyun Cao, Edward Courchaine, Rupak Datta, Giuseppe DeIuliis, Bertie Geng, Ryan Handoko, Chaney Kalinich, William Khoury-Hanold, Daniel Kim, Lynda Knaggs, Maxine Kuang, Joseph Lim, Melissa Linehan, Alice Lu-Culligan, Anjelica Martin, Irene Matos, David McDonald, Maksym Minasyan, M. Catherine Muenker, Nida Naushad, Allison Nelson, Jessica Nouws, Abeer Obaid, Annsea Park, Hong-Jai Park, Xiaohua Peng, Mary Petrone, Sarah Prophet, Tyler Rice, Kadi-Ann Rose, Lorenzo Sewanan, Lokesh Sharma, Denise Shepard, Michael Simonov, Mikhail Smolgovsky, Nicole Sonnert, Yvette Strong, Codruta Todeasa, Jordan Valdez, Sofia Velazquez, Pavithra Vijayakumar, Annie Watkins, Elizabeth B. White, Yexin Yang, Christina Harden, Maura Nakahata

